# Parasite-induced replacement of host microbiota- Impact of *Xenos gadagkari* parasitization on the microbiota of *Polistes wattii*

**DOI:** 10.1101/2024.11.26.625338

**Authors:** Deepak Nain, Anjali Rana, Rhitoban Raychoudhury, Ruchira Sen

## Abstract

**Background:** The study of microbiota of social insects under different ecological conditions can provide important insights into the role of microbes in their biology and behaviour. *Polistes* is one of the most widely distributed and extensively studied genera of social wasps, yet the microbiota of any species of *Polistes* or any primitively eusocial wasp is still unknown. *Polistes wattii* is an abundantly distributed Asian wasp which hibernates in winter and exhibits a unique biannual nest founding strategy. It is often parasitized by the strepsipteran endoparasite *Xenos gadagkari,* which changes the morpho-physiology and behaviour of their hosts. In this study we employ 16S rRNA amplicon sequencing, using the Oxford Nanopore platform, to study the microbial community associated with *P. wattii* and compared the microbiota of the unparasitized male and female *P. wattii* with their parasitized counterparts.

**Results:** The microbiota of females from preemergence solitary foundress nests differs from the females from the late colony phase of multiple foundress nests. The males and females also differ in their microbiota. To estimate the effect of strepsipteran parasitism on the microbiota of *P. wattii,* we compared the unparasitized wasps with that of the female and male *X. gadagkari* parasitoids and parasitized wasps. We show that the microbiota of the parasitoids and the parasitized wasps are dominated by *Wolbachia* and *Providencia.* β-diversity differences indicated that the parasitoid replaces and homogenizes the microbiota of *P. wattii.* Although the normal microbiota of *P. wattii* resembles that of highly eusocial vespid wasps, we show *X. gadagkari* parasitization replaces the microbiota of *P. wattii* and it becomes more like the microbiota of strepsipterans. Therefore, it appears that *X. gadagkari* and other such strepsipteran parasitoids may have a bigger impact on the biology of their hosts, than previously thought.

**Conclusion:** This is the first study of microbiota of any primitively eusocial Polistine wasp and any strepsipteran parasitoid associated with a hymenopteran species. Although strepsipteran parasitoids are commonly found in hymenopteran insects, the change on the microbiota of the host due to the parasitism was not known before. This study shows the drastic impact of the parasitoid on the microbiota of the host.

## Introduction

The association with microbes is an integral part of the biology of multicellular organisms (Rosenberg and Gophna 2011). Particularly, symbiotic association with microbes plays crucial roles in the survival, resource utilization, and defense against pathogens of many organisms including insects (Holt et al. 2024; Brownlie and Johnson 2009; Douglas 2015; A. Gupta and Nair 2020; Mondal et al. 2023; Kaltenpoth and Engl 2014). To estimate the presence of bacterial communities and their possible roles as symbiotes, the microbiota of selected body parts or whole bodies of insects have been studied (Suenami, Koto, and Miyazaki 2023; Suenami, Konishi Nobu, and Miyazaki 2019; Rothman et al. 2021; Cini et al. 2020; Chanson, Moreau, and Duplais 2023; Schwarz, Moran, and Evans 2016; Ishak, Miller, et al. 2011; Ishak, Plowes, et al. 2011; Otani et al. 2019; Shukla et al. 2016). Microbiota (particularly gut microbiota) of social insects have been shown to play crucial roles in the survival of their insect hosts. The effect of gut microbiota has been proposed in neurophysiology (Liberti and Engel 2020), nestmate recognition (Vernier et al. 2020), specific behaviour (Liberti et al. 2024) and also in the diet (Kešnerová et al. 2020) and longevity of honeybees (K. E. Anderson et al. 2018; Brown et al. 2022). The role of microbiota has also been linked with controlling weedy fungi and pathogenic fungi in fungus-growing ants (Sen et al. 2009) and termites (Agarwal et al. 2024; 2022). Microbiota differs in different developmental stages, populations and species of hornets (Zhang et al. 2022; Pang et al. 2023; Cini et al. 2020; Suenami, Konishi Nobu, and Miyazaki 2019). Thus, the association of highly eusocial insects (ants, honeybees, hornets and termites) with their resident microbes has been well established. However, the microbiota of primitively eusocial species remains relatively unexplored (Turillazzi, Meriggi, and Cavalieri 2023). Polistine wasps of the genera *Polistes*, *Ropalidia* and *Myschocyttarus* are standard model systems for many behavioural studies, which have been instrumental in understanding the nuances of sociality. However, they have been completely neglected in the field of microbial association studies.

*Polistes wattii* (Hymenoptera:Vespidae) is a widespread primitively eusocial wasp found in Asia and shows an unusual nest founding strategy. Single wasps initiate nests in spring, but multiple wasps initiate nests in summer. The spring nests are usually abandoned within 2-3 months before or after producing only a few wasps. The summer nests tend to last till the onset of winter and can expand to gigantic sizes, with multiple combs and produce males and female reproductives, destined for overwintering. This season-specific nest founding strategy is not known from any other primitively eusocial wasps (Sen et al. 2022; Nain and Sen 2023). Here we detail the pan- and core microbiota of *P. wattii* across seasons and tissues for both sexes. To our knowledge, this is the first detailed study on the microbiota of any Polistine or other primitively eusocial wasp species.

*P. wattii* wasps are often parasitized by an endoparasite *Xenos gadagkari* (Strepsiptera: Xenidae) (Nain et al. 2024). *Xenos* infections are reported in different species of *Polistes* and they cause severe behavioural and morpho-physiological changes in the host. The parasitized wasps leave their natal nest and join nest-free aggregations of other such parasitized wasps and subsequently become agents for spreading the infective stage of *Xenos* to new nests. (Hughes et al. 2004; Manfredini, Benati, and Beani 2010; Laura Beani and Massolo 2007; Laura Beani et al. 2011; Laura Beani 2006). These effects are so severe that Beani et al (Laura Beani et al. 2011) proposed to treat these infected wasps as a separate cast. Surprisingly, there are no reports of the microbiota of any species of *Xenos* or their impact on the microbial community of *Polistes*. To mitigate this gap, we have identified the pan- and core microbiota of both sexes of *X. gadagkari* and then identified how their parasitization changes *P. wattii* microbiota. Here we show that the infectious transfer of microbes from *X. gadagkari* to *P. wattii* leads to a drastic change in the host’s microbiota. This is the first study that investigates the effect of a strepsipteran parasitoid on the microbiota of any hymenopteran host. We further compared the microbiota of *P. wattii* with that of other highly eusocial hymenopterans (ants, bees and wasps). We conclude that the microbiota of unparasitized female *P. wattii* is similar to the highly eusocial wasps but parasitization influences the microbiota of parasitized wasps to shift away from their normal cluster.

## Materials and methods

### Wasp Collection, Dissection and DNA Extraction

*P. wattii* wasps were collected from freshly initiated nests in spring and late nests of summer from the Indian Institute of Science Education and Research (IISER), Mohali, campus (30.7046° N, 76.7179° E) (Supplementary Table 1) to sample the different castes. Wasps were kept in absolute ethanol till further dissection and processing. Unparasitized wasps (uninfected by *X. gadagkari*) were opened dorsally, and their guts were removed followed by DNA isolation of the gut as well as the rest of the body (carcass). The parasitized wasps (infected with *X. gadagkari*) were also opened similarly, but the parasitoids were carefully removed before extracting the gut tissue.

Before DNA extractions, all samples were surface sterilized with fresh ethanol (three rinses), followed by a rinse with autoclaved distilled water for tissue hydration. Gut and carcass tissues were then individually crushed with sterile micropestles in 200 µl of sterile Lysis Buffer (20 mM Tris [pH 8.0], 2 mM EDTA, 1.2% Triton X-100) and DNA was extracted using the PCI (Phenol: Chloroform: Isoamyl alcohol) method. Three biological replicates were used for each sample.

### Running PCR and Nanopore Platform for *16S* rRNA Amplicons

To identify the microbiota the V3-V4 region of the bacterial *16S* rRNA gene was amplified using primers 341F (5’-CCTAYGGGRBGCASCAG-3’) and 806R (5’-GGACTACNNGGGTAT CTAAT-3’) (Takahashi et al., 2014). Each 20 μl PCR reaction contained 14.5 μl of sterile water, 2 μl of 10X Buffer (Himedia) with 25mM MgCl2, 0.25 μM of each primer, 0.20 μM of dNTPs and 2 μl of template DNA. PCR cycles started with an initial denaturation at 95°C for 5 minutes, followed by 33 cycles at 95°C for 30 s, 56°C for 45 s, 72°C for 60 s, with a final extension at 72°C for 10 min.

Each biological replicate was further amplified in triplicates, and the PCR products of each biological replicate were pooled and purified using the QIAquick PCR Purification Kit (Qiagen). We followed the detailed protocol for library preparation as given in Agarwal et al. (2022). Briefly, purified PCR products were quantified using the Qubit dsDNA HS Assay Kit (Life Technologies) and were pooled in equimolar ratios, to approximately 300 ng DNA total. These were barcoded and 75 µl of the adapter-ligated pooled library was sequenced on a MinION platform (MinION Mk1B) on R9.4.1 MinION flow cell (Oxford Nanopore Technologies) using MinKNOW software with the protocol *NC_72Hr_sequencing_FLO-MIN106_SQK-LSK109_plus_Basecaller*. The MinION instrument was run for 72 hours until further sequencing reads could no longer be collected.

After completing the run, FASTQ files were generated by the *MinKNOW* software. Sequences were separated and trimmed according to their barcodes and DNA reads with Q > 7 were selected for further analysis with *Nanofilt* (De Coster et al. 2018). Various output metrics of these runs are summarized in Supplementary Table 2.

### Estimating Pan- and Core-Microbiota and Other Statistical Analysis

FASTQ files were converted to FASTA files and then the barcode and adaptor sequences were removed using the *Porechop* tool (Loman and Quinlan 2014). The sequences were then filtered by size using the *Seqtk* command, retaining 150–900 bp sequences for the V3-V4 region. Bacterial operational taxonomic units (OTUs) were identified using LAST v 1268 against a customized bacterial repository made by combining all the available *16S* rRNA gene sequences from the NCBI RefSeq 16S ribosomal RNA database (last accessed on 03^rd^ March 2023) and the Silva database (release 138) (Quast et al. 2013), with the following parameters: match score of 1, gap opening penalty of 1, and gap extension penalty of 1. Sequences with ≥80% identity were retained (confidence level filter) for further analysis. Sequences identified as fungal, archaeal, mitochondrial, or of chloroplast origin were removed. To estimate the sequencing depth, rarefaction curves were generated with the Vegan package (Oksansen et al. 2013) v 2.5–4 in R v 4.1.3. Taxonomic richness, diversity, and evenness were also determined in R using the nonparametric species richness estimator Chao 1, Shannon, and Simpson indices. To account for sequencing depth differences, samples were rarefied and normalized to the sample size with the lowest number of reads (449 sequences obtained from Unparasitized Male Carcass) to ensure a random subset of bacterial OTUs for all samples (averaged across10 random replicates). These rarefied bacterial OTUs were used to calculate the different α-diversity indices. Pan-microbiota was established by enumerating all the bacterial OTUs found from carcass and gut for *P. wattii*. Bacterial OTUs common for any set of samples were designated as core-microbiota. For both these measures bacterial OTUs ≥ 0.3% of relative abundance was used for each sample. The pan- and core microbiota were visualized using a Venn diagram (https://bioinformatics.psb.ugent.be/webtools/Venn). Chord diagrams were also generated for the core microbiota using the *circlize* package in R v4.1.3.

Weighted UniFrac distances (Lozupone and Knight 2005) were used to evaluate the β- diversity that accounts for phylogenetic relationships and the abundance of bacterial OTUs. Principal coordinate analysis (PCoA) was used to visualize distance matrices using the *Phyloseq* package (McMurdie and Holmes 2013) in R. Bray–Curtis Dissimilarity analysis (Oksansen et al. 2013) was also used to compare the differences between microbiota samples based on the abundance of bacterial OTUs. Dissimilarity matrices between samples were generated and visualized through Hierarchical Cluster Analysis (HCA) using the *Phyloseq* package in R. The index ranges from 0 to 1, with 0 indicating nearly equal representation of all bacterial OTUs and 1 indicating unequal representation (abundance). To statistically test consistent compositional and abundance differences in the different microbiotas the Permutational Multivariate Analysis of Variance (PERMANOVA) (M. J. Anderson 2017) was used. These were performed through the *vegan* package in R v 4.1.3, where each comparison was performed with 999 permutations. The heatmap for all the samples was generated using the *pheatmap* package in R v4.1.3. High-quality sequences were deposited to NCBI Sequence Read Archive (SRA) under the BioProject PRJNA1182662, the microbiota of *P. wattii* samples with the BioSample accessions SAMN44601033-SAMN44601043 and the microbiota of *X. gadagkari* samples with SAMN44601044 and SAMN44601045.

### Multi Locus Strain Typing (MLST) of *Wolbachia* Infection in *X. gadagkari* and Parasitized *P. wattii*

*Wolbachia* reads were detected in *X. gadagkari* and parasitized *P. wattii* samples from Nanopore sequencing data. The presence of *Wolbachia* was further confirmed by successful amplification of the *16S* rRNA region of *Wolbachia* using Wspec-F and Wspec-R primers (Werren and Windsor 2000) (Supplementary Table 3). The characterization of the *Wolbachia* strain was performed by sequencing multiple loci (gatB, coxA, hcpA, ftsZ, and fbpA) recommended by the *Wolbachia* MLST system (http://pubmlst.org/Wolbachia) (Baldo et al. 2006) (Supplementary Table 3). To test whether uninfected tissue samples were contaminated by the presence of cryptic *X. gadagkari*, we amplified and sequenced the mitochondrial *CO1* region with the cox1F/cox1R primers (Supplementary Table 3). The chromatograms of the sequences obtained were screened for the presence of *X. gadagkari* sequence (NCBI accession OR086007-OR086008) by looking for corresponding multiple peaks in Sequencher (Gene Codes Corporation, USA). Sequences which only showed unambiguous *P. wattii CO1* peaks (NCBI accession MT891254-MT891261) were used as uninfected samples and were further used for Nanopore sequencing.

### Estimation of the relative density of *Wolbachia* infection through qPCR Assay

To assess the infection dynamics of *Wolbachia* in parasitized *P. wattii* and *X. gadagkari*, different body parts of *P. wattii* were probed for the presence of *Wolbachia*. Gut from unparasitized male and solitary foundress was used as negative controls to further confirm the absence of *Wolbachia* in unparasitized *P. wattii*. qPCR was performed for three biological replicates of each sample using the CFX96 C1000® Touch Real-time qRT-PCR machine (BioRad). The amplification was carried out for the *Wolbachia hcpA* gene (Tiwary et al. 2022) (Supplementary Table 3). The heat shock protein (*HSP*) gene of both the host and the parasitoid was utilized as the control gene (Jandt et al. 2015) (Supplementary Table 3). The qPCR reactions were performed in a total volume of 10 μl, containing 5 μl of iTaq Universal SYBR® Green supermix (BIORAD), 0.05 μl each of 10 μM of forward and reverse primers, and 100 ng of template DNA. *Wolbachia*-uninfected *Nasonia vitripennis* DNA was used as another negative control, while DNase-free water was used as a no-template control. The reaction conditions included an initial denaturation step of 95°C for 3 minutes, followed by 39 cycles of 95°C for 10 s, annealing at 54°C for 30 s, and amplification at 72°C for 10 s. All the reactions were performed in triplicates and included a melt curve analysis to check for non-specific amplification. The relative *Wolbachia* density was estimated by calculating the mean delta threshold cycle (ΔCt), where we normalized the Ct values of *Wolbachia* to the Ct values of the housekeeping gene (*HSP* gene) to correct for the variations between samples. The relative quantifications (RQ) were calculated and plotted to show the *Wolbachia* density in different *P. wattii* and *X. gadagkari* samples using the formula (Livak and Schmittgen 2001).

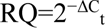

All the results were statistically analyzed using R software (R v4.1.3) through one-way ANOVA followed by a multiple comparison test (Tukey’s post-hoc test) with a significance level of 0.05%. Primer efficiency was also checked through a standard curve analysis based on 10-fold serial dilutions of purified larger PCR product of *Wolbachia hcpA* gene of known concentrations.

## Results

### Microbiota of *P. wattii* is different across seasons and sex

The Nanopore runs yielded over 1.1 x 10^6^ high-quality reads (Supplementary Table 2) across all the samples tested. These were identified to 535 bacterial OTUs (Supplementary Table 2). The bacterial diversity with their relative abundances is presented in Figure 1 across nesting habit, sex, infection status and developmental stage (Figure 1) and in a heatmap (Supplementary Figure 1). The rarefaction curves for most samples tended towards saturation, indicating adequate sequencing (Supplementary Figure 2). The α-diversity indices of each sample are given in Supplementary Table 2.

**Figure 1:**
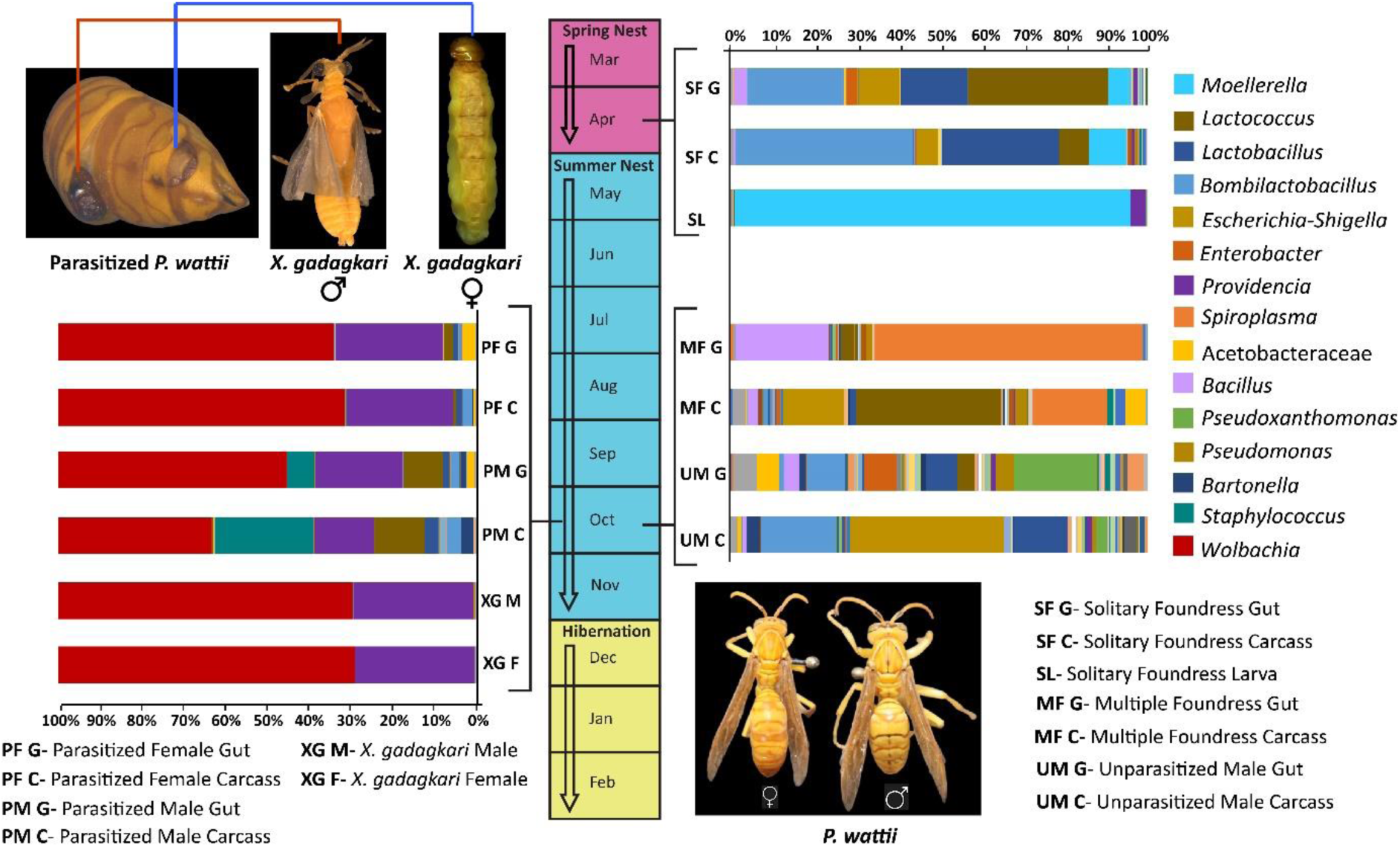
**Microbiota of *Polistes wattii* and *X. gadagkari***. The central scale represents the annual nesting cycle of *P. wattii.* The right panel demonstrates the relative abundance of bacteria at genus level in unparasitized *P. wattii* based on season and sex. The left panel demonstrates the relative abundance of bacteria at genus level of parasitized male and female *P. wattii* and male and female *X. gadagkari*. Each bar represents the proportions of different bacterial OTUs (relative abundance expressed as a percentage). The plot highlights bacterial genera with an abundance of 0.3% or greater.

*P. wattii* uses haplometrosis or solitary founding (SF) strategy in spring and switches to pleiometrosis or multiple founding (MF) strategy in summer. Therefore, to have a comprehensive understanding of the microbiota of *P. wattii* females, we analyzed the gut and carcass of unparasitized females from SF nests, as well from unparasitized females from MF nests (Supplementary Table 1 and Figure 1). The pan-microbiota of unparasitized *P. wattii* females revealed 62 bacterial OTUs (Figure 1) whereas, the core microbiota consisted of 22 unique OTUs (Figure 2). Although the SF and MF wasps had 46 shared bacterial genera, these seasonal samples differed as *Lactobacillus*, *Bombilactobacillus* and *Moellerella* were more abundant in SF females while *Bacillus, Spiroplasma* and *Pseudomonas* were more abundant in MF females. β-diversity analysis indicated significant differences between the microbiota of SF and MF females, both in their pan-microbiota (pairwise PERMANOVA, *p* = 0.006; Supplementary Table 4) as well as their core microbiota (pairwise PERMANOVA, *p* = 0.015; Supplementary Table 5). γ- proteobacteria dominated the microbiota of solitary foundress larvae, with *Moellerella* representing almost the entirety of the microbiota (93.3% relative abundance), indicating a much lower diversity. *Moellerella* was also detected in the gut and carcass (5.38% and 8.44%, respectively), of the SF and was not detected in adult males and females from MF nests.

**Figure 2:**
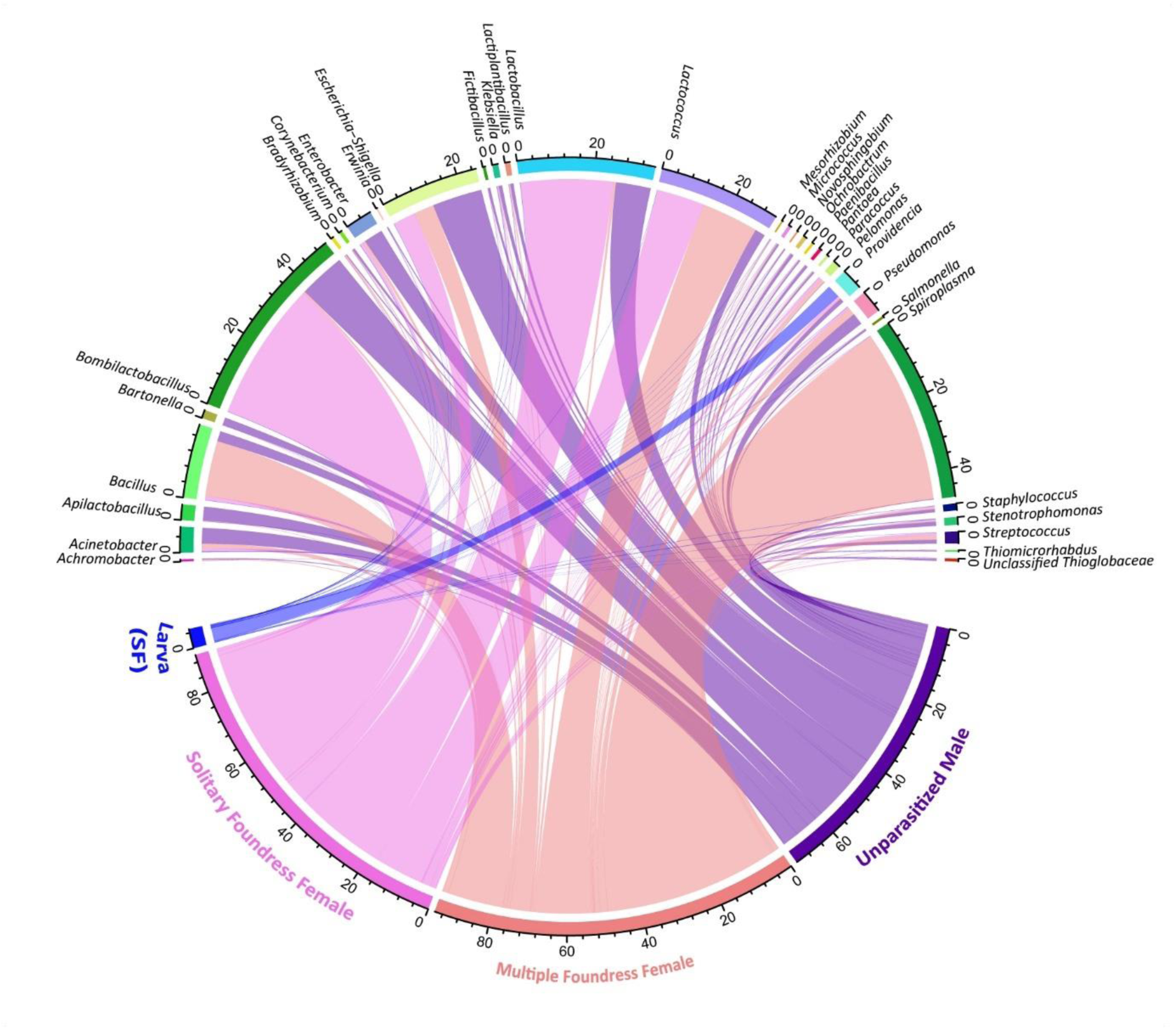
Prevalence of the shared bacterial OTUs in unparasitized *P. wattii.* Females from Solitary and multiple foundress nests were collected in April and October respectively. Males were collected from multiple foundress nests in October.

The pan-microbiota of unparasitized *P. wattii* males revealed 62 bacterial OTUs (Figure 1) whereas, the core-microbiota consisted of 33 unique OTUs. β-diversity of the core-microbiota of the males differed significantly with SF females (pairwise PERMANOVA, *p* = 0.013; Supplementary Table 5), as well as MF females (pairwise PERMANOVA, *p* = 0.011; Supplementary Table 5). Similar differences were also found for the pan-microbiota (pairwise PERMANOVA, *p* = 0.009, *p* = 0.005 for SF females and MF females, respectively; Supplementary Table 4). 16 shared bacterial OTUs were found between the sexes (Supplementary Figure 3a). However, *Pseudoxanthomonus* and *Xanthomonus* dominated the microbiota of males while *Lactococcu*s, *Lactobacillus, Bacillus,* and *Spiroplasma* were more abundant in females (Figure 1, Supplementary Table 6).

### Microbiota of *Xenos gadagkari* shows limited diversity

The microbiota of the male and female *X. gadagkari* were similar as 17 out of 31 OTUs were shared across them with nearly equal abundances. Moreover, 8 out of these 17 core bacteria were also found to be common with the core microbiota of unparasitized *P. wattii* (Supplementary Figure 3b). This shows a limited diversity of the *X. gadagkari* microbiota with only 9 different bacteria unique to it. The two most abundant bacterial OTUs found were from *Wolbachia* and *Providencia* with ∼70% abundance and 28% abundance, respectively.

### *X. gadagkari* parasitization replaces the *P. wattii* microbiota

We found a significant impact of parasitization by *X. gadagkari* on the composition of *P. wattii* microbiota. The microbiota of the parasitized wasps mainly differed from the unparasitized ones by the increased abundance of *Wolbachia* and *Providencia* and the reduced abundance of the core bacteria of *P. wattii*, including, *Bacillus, Escherichia-Shigella*, *Pseudomonas, Spiroplasma, Pseudoxanthomonas* and *Spiroplasma* (Figure 1). *Wolbachia* and *Providencia* were prevalent in the gut and carcass of both male and female parasitized wasps, indicating transfer from parasitoid to host (Figure 1). This overlap is described in Figure 3.

**Figure 3:**
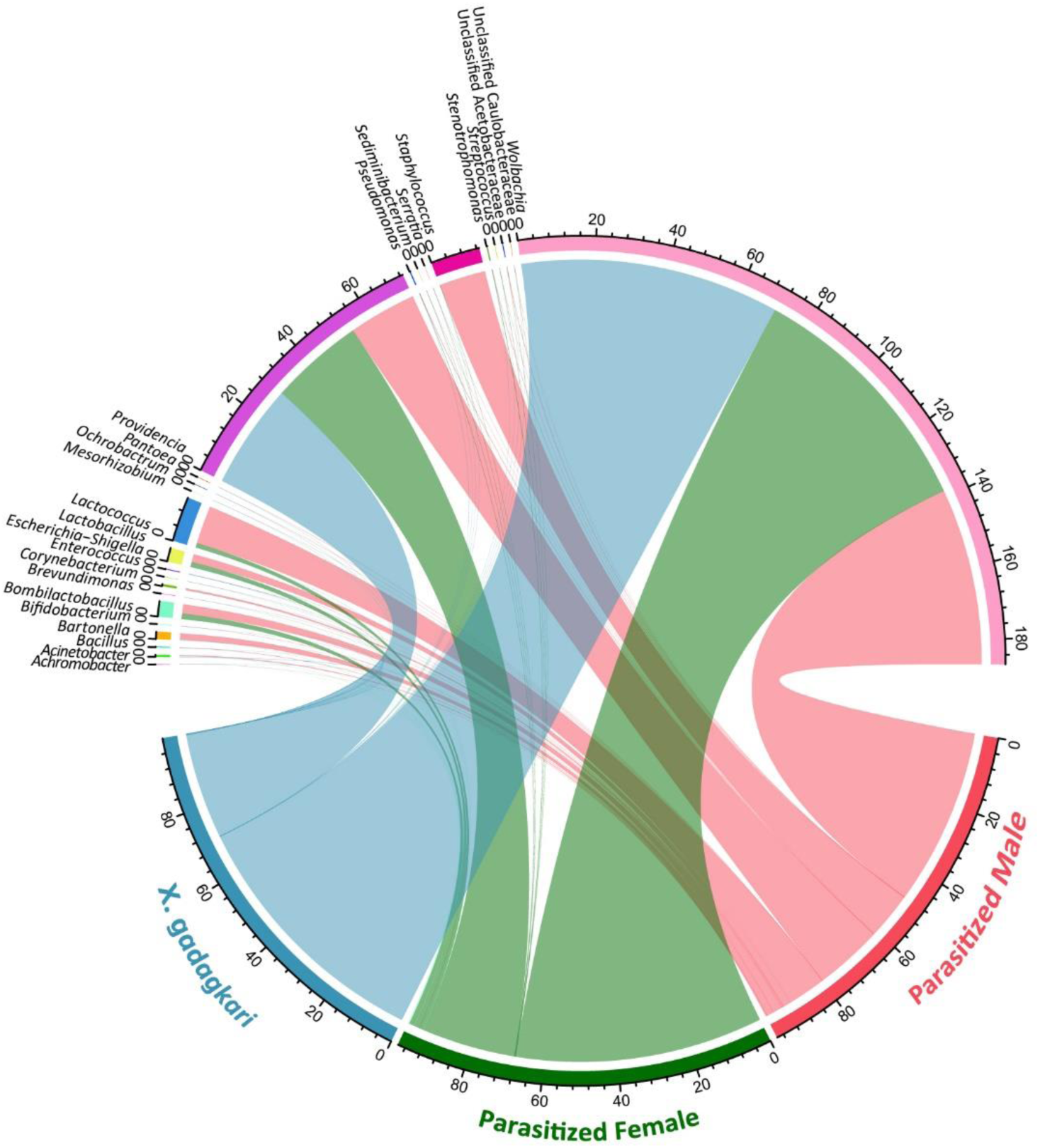
Prevalence of the shared bacterial OTUs between parasitized *P. wattii* samples and *X. gadagkari* samples.

Principal coordinate analysis (PCoA) based on weighted-Unifrac distances was carried out (Figure 4) to compare the microbiota of unparasitized and parasitized wasps and revealed a clear separation into two groups (PERMANOVA, *p* = 0.001). All samples of unparasitized *P. wattii* clustered together, regardless of tissue type, sex, and developmental stage. The parasitized *P. wattii* samples clustered with *X. gadagkari* samples, suggesting a greater degree of similarity between the parasitoid and the host.

**Figure 4:**
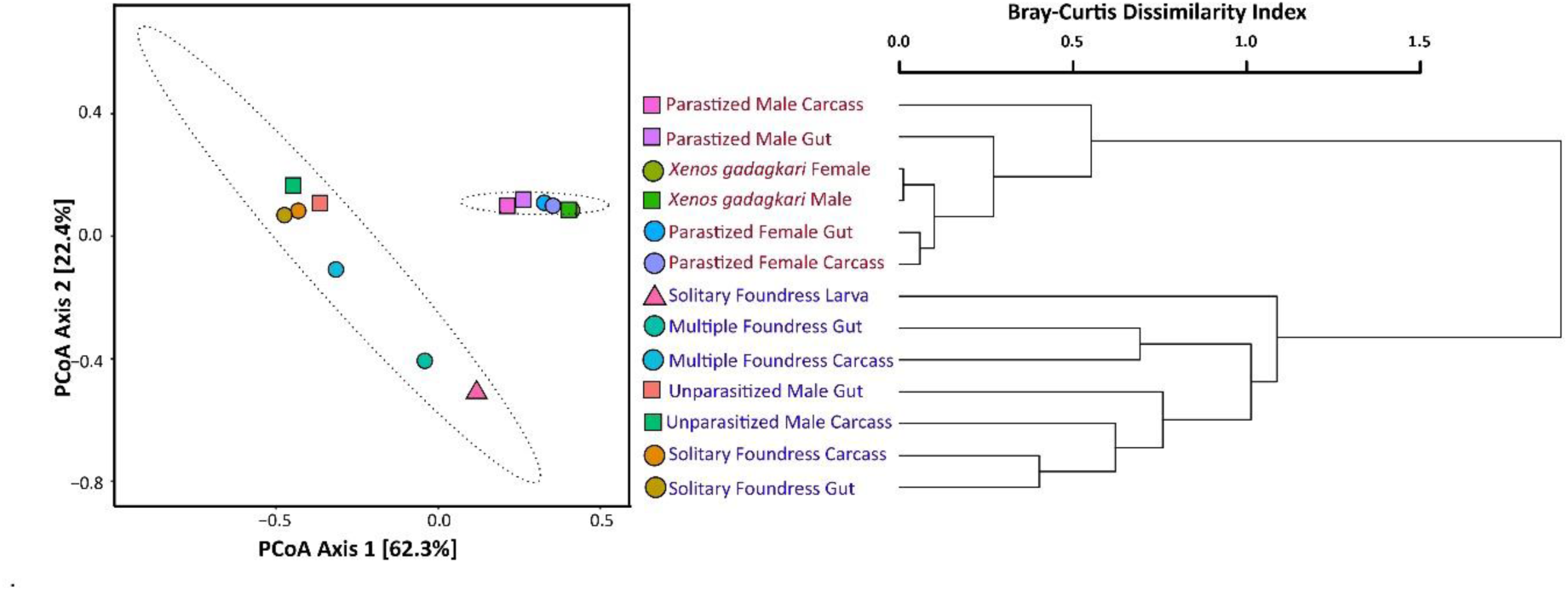
β**-diversity analysis.** A) Principal Coordinate Analysis (PCoA) using Weighted UniFrac depicting β diversity across different *P. wattii* and *X. gadagkari* samples. Color represents different samples. Shape shows sex or developmental stage. (B) Hierarchical Cluster Analysis (HCA) of *P. wattii* and *X. gadagkari* samples based on Bray-curtis dissimilarity matrix.

These results demonstrate that *X. gadagkari* parasitization is a major factor in determining the microbiota composition of *P. wattii*, with the first two principal coordinates (PCo1 and PCo2) together explaining 84.70% of the variation in the data. This was further supported by hierarchical cluster analysis (HCA), based on the Bray-Curtis dissimilarity matrix (Figure 4), which showed clustering of parasitized *P. wattii* with *X. gadagkari*. All unparasitized *P. wattii* samples formed a separate clade, including the solitary foundress larva, showing a greater effect of parasitization on the microbiota of *P. wattii* than developmental stages. HCA also showed separate clades for SF and MF nest samples indicating that the microbiota composition differs across seasons. However, the exception to this seasonal clustering were the male samples which formed a clade with the SF samples, rather than MF nests, despite being collected from the MF nests.

### Introduction of *Wolbachia* Infections into *P. wattii* by *X. gadagkari* parasitization

The microbiota of *X. gadagkari* was less diverse than the microbiota of their hosts (Figure 1). Certain bacteria (like *Apilactobacillus, Lactobacillus*, *Bombilactobacillus, Lactococcus, Escherichia-Shigella and Paracoccus*) that are present in high abundance in the hosts are either not present in *X. gadagkari* or are represented by only a few sequences (less than 0.3% abundance). *Paenibacillus,* was commonly found in wasp samples albeit in low abundance but it could not be detected in *X. gadagkari*. Instead, it was dominated by the α-proteobacteria, *Wolbachia* (approximately 70% abundance), and γ- proteobacteria, *Providencia* (more than 28% abundance) (Figure 1).

Infections of the endosymbiont *Wolbachia* was found in *X. gadagkari* and parasitized, but not unparasitized, *P. wattii* samples (Figure 1). This indicates that *X. gadagkari* parasitization is responsible for the introduction of *Wolbachia* into parasitized *P. wattii*. Screening of *P. wattii* samples consistently gave *Wolbachia*-positive signals for parasitized wasps but never for unparasitized ones. The Multi Locus Strain Typing (MLST) system for *Wolbachia* (Baldo et al. 2006) revealed the *X. gadagkari Wolbachia* consisted of previously identified alleles (coxA-23, ftsZ-3 (closest match with 1bp difference), gatB-22, fbpA-23, and hcpA-24) establishing it as A-supergroup *Wolbachia* infection (Baldo et al. 2006). Moreover, the sequence of *ftsZ* gene of *Wolbachia*, from parasitized *P. wattii* and *X. gadagkari*, was also found to be identical. Sequences of gatB, ftsZ, fbpA, coxA and hcpA genes of *X. gadagkari* have been deposited in the NCBI GenBank database under accession numbers PQ635193-PQ635197. Sequence of the ftsZ gene of parasitized *P. watti* was submitted under accession number PQ635192. To further analyze whether these *Wolbachia* infections are consistently transferred to *P. wattii* we determined their densities through qPCR, which revealed significant variations in the density of *Wolbachia* among the parasitized *P. wattii* and *X. gadagkari* samples (Supplementary Figure 4). Comparison of relative quantification, or fold change, of *Wolbachia* in parasitized male and female *P. wattii* with male and female *Xenos* showed significant differences (One-way ANOVA p= 0.0024, F= 4.52, df= 9). *X. gadagkari* females showed the highest density of *Wolbachia* infections (Supplementary Table 7, Supplementary Figure 4) indicating that this is the most likely source of this *Wolbachia* strain.

## Discussion

### Microbiota of *P. wattii* shows seasonal and sex bias

This is the first detailed study on the microbiota of any primitively eusocial wasp like *Polistes*. With a species richness of over 200, and distribution in most parts of the world, *Polistes* shows one of the most significant adaptive radiations of primitively eusocial wasps. Due to their small colony size and open nesting habit, many species of *Polistes* are extensively used as model systems for behavioural studies (Reeve 1991)., In comparison to temperate and neotropical *Polistes, P. wattii* further shows at least two unique features- the alternative biannual nesting strategy and the construction of massive nests with multiple combs in summer (Sen et al. 2022). The alternation of haplometrosis (SF) strategy in spring and pleiometrosis (MF) strategy in summer (detailed in Figure 1) should influence the microbiota of these wasps. *P. wattii* reproductive females and males are usually produced at the end of the colony cycle (Oct-Nov), where females mate and undergo reproductive diapause till the beginning of spring (March). Our data indicates that there is a significant difference in gut microbiota of males and females of MF nests, despite being collected from the same season around the same time and, presumably, being fed similar seasonally available food (Figure 1, pairwise PERMANOVA, *p* = 0.005; Supplementary Table 4). *P. wattii* emerges from diapause in March and April and initiate fresh solitary nests but most of these are abandoned within a few months (Sen et al. 2022). This seasonal variation between the SF and MF females also turns out to be correlated with the significant difference in their microbiota (pairwise PERMANOVA, *p* = 0.006; Supplementary Table 4; Figure 1). What roles, if any, these seasonally different microbes play in *Polistes* biology need further investigation, especially comparisons with other primitively eusocial hymenopterans.

### Replacement of *P. wattii* microbiota due to *X. gadagkari* parasitization

Strepsipteran parasitoids infect a wide range of insects (Kathirithamby 2009) and their drastic effects on the morpho-physiology and behaviour of the polistine hosts are already well established (Laura Beani et al. 2011; Laura Beani and Massolo 2007; Manfredini, Benati, and Beani 2010; Hughes et al. 2004). However, the impact of strepsipteran parasitoids on the host microbiota remained unexplored. The microbiota of the strepsipteran parasitoid has been studied in *Dipterophagus daci* (Family Halictophagidae), which is a parasite of tephiritid fruitflies (Towett-Kirui, Morrow, and Riegler 2022; Towett-Kirui et al. 2023). Like *P. wattii* and *X. gadagkari*, the microbiota of *D. daci* was also found to be distinct (dominated by *Wolbachia*) and less diverse than its hosts*. X. gadagkari*, being a parasitoid, subsists on *P. wattii*, and therefore, some bacterial exchange is inevitable. This is reflected in the similarities in α-diversity estimates of parasitized females and *X. gadagkari* microbiota (Supplementary Table 2, Figure 1 and Supplementary Figure 2). However, if similar bacteria are transferred from *X. gadagkari* to *P. wattii*, then the relative abundances of these transferred bacteria would be higher in infected *P. wattii* than their unparasitized counterparts. This can be better captured by β-diversity estimates which is influenced more by abundance than by the presence of unique bacterial OTUs (Lozupone and Knight 2005; Oksansen et al. 2013). Accordingly, we found the relative abundances of *Providencia, Staphylococcus* and *Wolbachia* to have substantially increased in the parasitized wasps (Figure 1 and Supplementary Table 6). Another possible outcome of such an exchange is the homogenization of the microbiota of the parasitized host with the parasitoid (Gloder et al. 2021). In *P. wattii*, the gut microbiota of unparasitized and parasitized females are 4.66% similar (Bray-Curtis Dissimilarity Analysis; Supplementary Table 8). Moreover, post-parasitization, *P. wattii* gut microbiota becomes 90.06% similar to *X. gadagkari*. This is also true for the carcass microbiota, where the corresponding similarities increase from 3.41% to 78% similarity (Figure 4 and Supplementary Table 8). However, there is a sex bias as male *P. wattii* is not homogenized to the same extent as the female microbiota. Post-parasitization by *X. gadagkari*, the microbiota of male gut becomes 61.31% similar to *X. gadagkari* (Figure 4 and Supplementary Table 8).

### Introduction of *Wolbachia* through strepsipteran parasitization

The dynamics of *Wolbachia* infections, introduced to *P. wattii* by *X. gadagkari* parasitization, also provides evidence for microbiota replacement. *Wolbachia* is a widely prevalent maternally inherited endosymbiont of arthropods, which can elicit several reproductive alterations on its host by acting as a selfish bacteria. *Wolbachia* spreads across species largely by horizontal transfer, where it is transferred from one host to another taxonomically unrelated one. We could not detect *Wolbachia* infections in unparasitized *P. wattii,* but it was consistently detected in *X. gadagkari* and parasitized *P. wattii*. This means the source of *Wolbachia* is *X. gadagkari.* Further evidence for this comes from qPCR studies on different tissues of parasitized *P. wattii* and *X. gadagkari* itself, where the highest density of infection was always found in the parasitoid and not the host. This pattern of infection, *i.e*., the presence of *Wolbachia* in parasitized but not in unparasitized fly hosts, has also been seen in the strepsipteran *D. daci* (Towett-Kirui et al. 2023). Such transfer of *Wolbachia* through parasitoids is not uncommon (Noda et al. 2001; Gloder et al. 2021; Towett-Kirui et al. 2023) and has been reported from several such systems of host-parasitoids and is one of the major mechanisms of transfer of *Wolbachia* across arthropod communities (M. Gupta et al. 2021). However, *Wolbachia* infection in *P. wattii* and *D. daci* has not established itself in the host population. This could be due to the effective castration of the female wasps by their strepsipteran parasitoids. Although male parasitized wasps are usually not castrated (L. Beani et al. 2017) they cannot transfer the maternally inherited *Wolbachia.* The widespread presence of *Wolbachia* across the parasitized hosts, including its head, can be indicative of the role of this bacteria in the morphophysiological and behavioural changes of the host. However, this remains to be investigated.

To understand the nature of the *P. wattii* microbiota, we compared the β-diversity of bacterial OTUs of female *P. wattii* with two highly eusocial wasps (*Vespa velutina* and *Vespula pensylvanica*) (Cini et al. 2020; Rothman et al. 2021), two ants (*Formica exsecta* and *F. lemani*) (Jackson et al. 2023) and the European honeybee (*Apis mellifera*) (Tola et al. 2020) by weighted-Unifrac PCoA (Figure 5). To see the effect of the parasitization, we also included the microbiota of parasitized female wasps, male and female *X. gadagkari* and male *D. daci* in this analysis.

**Figure 5:**
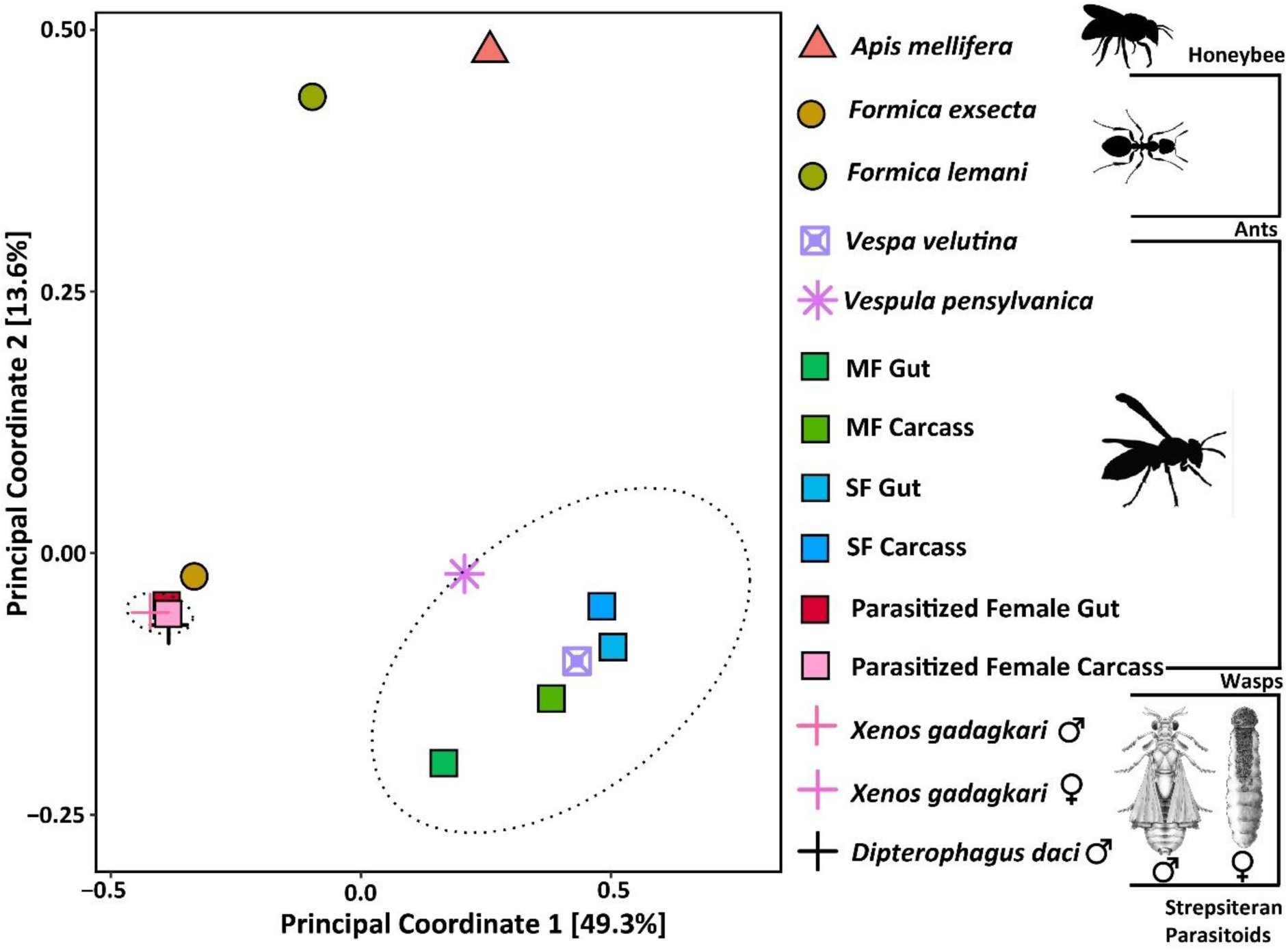
Comparison of the microbiota of *P. wattii* females and *X. gadagkari* with other hymenopteran social insects (Supplementary Table 9) and one strepsipteran parasitoid using the Principal Coordinates Analysis (PCoA).

The unparasitized *P. wattii* clustered with other wasps, away from the ants and the honeybee, indicating that the members of the Vespidae family may have similar microbiota (Figure 5, PERMANOVA, *p* = 0.001). This similarity could be a result of their similar carnivorous food habit. The impact of parasitization was prominent as the strepsipteran parasitoids (*X. gadagkari* and *D. daci*) and the parasitized wasps clustered together away from other unparasitized insects. The clustering of parasitoids and parasitized wasps was the result of the abundant presence of *Wolbachia* and the homogenization of the bacterial community within them. The microbiota of the unparasitized ant *F. exsecta,* which was also dominated by *Wolbachia,* was found close to this cluster. This could be indicative of a major pathway of *Wolbachia* spread across arthropod communities through their strepsipteran parasitoids.

What roles microbes play in the biology and colony cycle of social wasps remains unknown due to the lack of information on the microbiota of other primitively eusocial wasps. Which ecological pressure is responsible for *P. wattii* abandoning a high percentage of solitary nests in spring and requiring new nests in summer is also not known. This study of the detailed microbiota of *P. wattii*, from two different seasons across two different nesting strategies and parasitic status, will help us understand how different ecological factors impact the microbiota of this primitively eusocial species. Furthermore, a drastic change takes place in the behaviour of the *Xenos*-parasitized wasps, as they become unsocial and form aggregations with other such parasitized wasps. As a result of these morpho-physiological and behavioural changes brought about by *Xenos* parasitism, the female wasps can neither reproduce nor become workers. How *Xenos* overrules both these fitness options for *P. wattii* is not clearly understood. In this study we also show that the microbiota of these parasitized wasps is homogenized and replaced by *Xenos* infections. Whether these microbes play any part in this behavioural change of *P. wattii* remains to be investigated.

## Conclusions

The effects of season, sex and parasitism can be profound on microbiota of social insects. Here we show the microbiota of the primitively eusocial polistine wasp, *P. wattii* shows a seasonal variation, in accordance with its two different nesting strategies and sex. We further demonstrate that *P. wattii*, upon parasitization by *X. gadagkari*, have their normal microbiota replaced by this parasitoid. This replacement is particularly significant for the introduction of *Wolbachia* into *P. wattii*. We show that the presence of *Wolbachia* in different tissues of parasitized *P. wattii* but not in unparasitized wasps. This evidence adds to a few other cases of such transfer by strepsipteran parasitoid, indicating that *Wolbachia* can use this route to transfer into various host populations. However, since the parasitized wasps do not reproduce, the implication of transfer of *Wolbachia* is not clear. This is the first exploration of microbiota for any primitively eusocial wasp species and any strepsipteran parasitoid associated with a hymenopteran social insect. Whether our results are representative of other such primitively eusocial wasps, however, awaits further such studies.

## Declarations

### Ethics Approval and consent to participate

Not Applicable

### Consent for Publication

All authors agree to the submission of the manuscript and corresponding authors Ruchira Sen and Rhitoban Raychoudhury have been authorized by co-authors.

### Availability of data and material

All sequence data are submitted to Genbank. Accession numbers are given in the manuscript wherever applicable.

### Author contribution

RS and RR conceived and designed the study. DN and RS collected wasps. DN dissected the wasps. DN and AR carried out the molecular preparation and visualization, AR with DN conducted the Bioinformatics and statistical analysis under the supervision of RS and RR. The first draft of the paper was written and subsequently reviewed and edited by all authors.

## Acknowledgements

The authors are thankful to the Department of Forests and Wildlife, Chandigarh Administration and the Department of Forests and Wildlife Preservation, Punjab, for the permit and NOC to collect wasps.

