## Supplementary file for "Parasite-induced replacement of host microbiota- Impact of *Xenos gadagkari* parasitization on the microbiota of *Polistes wattii*"

**Title:**

**Supplementary Table 1.** Collection details of samples used for the preparation of Nanopore library of *Polistes wattii* and *Xenos gadagkari.*

| Sample | Collection Date | Collection Location in IISER Mohali | No. of identified bacterial reads |
|---|---|---|---|
| Solitary foundress female gut | 3 nests, 21-03-2023 | Academic Block 1 | 1,040 |
| Solitary foundress female Carcass | 3 nests, 21-03-2023 | Academic Block 1 | 3,636 |
| Solitary foundress Larva | 1 nest, 21-03-2023 | Academic Block 1 | 17,701 |
| Multiple foundress female gut | 18-10-2022<br>24-11-2021 | Nest 1- MJ Block<br>Nest-2-Visitors Hostel | 5,167 |
| Multiple foundress female Carcass | 18-10-2022<br>24-11-2021 | MJ Block,<br>Visitors Hostel | 3,340 |
| Unparasitized Male gut | 18-10-2022<br>24-11-2021 | MJ Block<br>Visitors Hostel | 908 |
| Unparasitized Male Carcass | 18-10-2022<br>24-11-2021 | MJ Block<br>Visitors Hostel | 449 |
| Parasitized Male gut | 18-10-2022<br>24-11-2021 | MJ Block<br>Visitors Hostel | 5,199 |
| Parasitized Male Carcass | 18-10-2022<br>24-11-2021 | MJ Block<br>Visitors Hostel | 1,071 |
| Parasitized Female gut | 18-10-2022<br>24-11-2021 | MJ Block<br>Visitors Hostel | 9,191 |
| Parasitized Female Carcass | 18-10-2022<br>24-11-2021 | MJ Block<br>Visitors Hostel | 14,666 |
| *Xenos gadagkari* Male | 24-11-2021 | MJ Block | 12,809 |
| *Xenos gadagkari* Female | 24-11-2021 | MJ Block | 9,523 |

**Supplementary Table 2:** Number of reads obtained to identify the microbiota of *P. wattii* and its parasite, *X. gadagkari*. The table also shows the diversity indices for all the samples.

| Sr No | Sample | Total reads obtained | High-quality reads | Bacterial reads identified | Bacterial OTUs identified | Chao1 | Shannon | Simpson |
|---|---|---|---|---|---|---|---|---|
| 1 | Solitary Foundress Gut | 8,497 | 3,207 | 1,040 | 48 | 91.875 | 2.0313899 | 0.799445 |
| 2 | Solitary Foundress Carcass | 18,966 | 5,866 | 3,636 | 132 | 386.75 | 2.0061795 | 0.747096 |
| 3 | Solitary Foundress Larva | 64,000 | 20,217 | 17,701 | 165 | 325 | 0.430308 | 0.128157 |
| 4 | Multiple Foundress Gut | 15,872 | 6,788 | 5,167 | 121 | 210.4375 | 1.4887225 | 0.563903 |
| 5 | Multiple Foundress Carcass | 38,092 | 6,334 | 3,340 | 254 | 447 | 3.0152656 | 0.857085 |
| 6 | Unparasitized Male Gut | 13,537 | 3,906 | 908 | 115 | 184.8333 | 3.5502066 | 0.939134 |
| 7 | Unparasitized Male Carcass | 12,612 | 3,965 | 449 | 67 | 120.0909 | 2.651658 | 0.83197 |
| 8 | Parasitized Female Gut | 20,313 | 9,460 | 9,191 | 94 | 175.0588 | 1.0805678 | 0.507146 |
| 9 | Parasitized Female Carcass | 36,000 | 21,896 | 14,666 | 136 | 234.7727 | 0.9979121 | 0.4761 |
| 10 | Parasitized Male Gut | 13,551 | 5,679 | 5,199 | 76 | 173 | 1.5284611 | 0.649796 |
| 11 | Parasitized Male Carcass | 11,074 | 4,269 | 1,071 | 64 | 175.4286 | 1.8766825 | 0.777282 |
| 12 | *X. gadagkari* Male | 23,920 | 13,260 | 12,809 | 87 | 153.5625 | 0.7425438 | 0.430457 |
| 13 | *X. gadagkari* Female | 16,697 | 9,561 | 9,523 | 61 | 126 | 0.6910322 | 0.422064 |
| | Total | **2,93,131** | **1,14,408** | **84,700** | | | | |

**Supplementary Table 3:** List of primers used in this study.

| Gene | Primer | Primer Sequence (5'-3') | Annealing Temp (˚C) | Amplicon Size (bp) | Reference |
|---|---|---|---|---|---|
| Cox1 | Cox1 22F | TCWACAAATCATAAAATAATTGG | 50 | 647 | Benda et al., 2021 |
| | Cox1 669R | TCCTCCTCCTAAAGGRTCRAA | | | |
| wspec | 16S wspecF | CATACCTATTCGAAGGGATAG | 55 | 500 | Werren and Windsor (2000) |
| | 16S wspecR | AGCTTCGAGTGAAACCAATTC | | | |
| gatB | gatB general F | GAKTTAAAYCGYGCAGGBGTT | 57.5 | 369 | Baldo et al. (2006) |
| | gatB general R | TGGYAAYTCRGGYAAAGATGA | | | |
| coxA | coxA general F | TTGGRGCRATYAACTTTATAG | 51 | 402 | Baldo et al. (2006) |
| | coxA general R | CTAAAGACTTTKACRCCAGT | | | |
| hcpA | hcpA general F | GAAATARCAGTTGCTGCAAA | 51 | 444 | Baldo et al. (2006) |
| | hcpA general R | GAAAGTYRAGCAAGYTCTG | | | |
| ftsZ | ftsZ general F | 5'-ATYATGGARCATATAAARGATAG | 54 | 435 | Baldo et al. (2006) |
| | ftsZ general R | TCRAGYAATGGATTRGATAT | | | |
| fbpA | fbpA general F | GCTGCTCCRCTTGGYWTGAT | 56 | 429 | Baldo et al. (2006) |
| | fbpA general R | CCRCCAGARAAAAYYACTATTC | | | |
| hcpA | hcpa_qPCR_F | CTTCGCTCTGCTATATTTGCTGC | 54 | 180 | Tiwary et al 2022 |
| | hcpa_qPCR_R | CGAATAATCGCAACCGAACTG | | | |
| HSP 90A | HSP 90A F | CGGCAGAGGATTCCCATAAA | 54 | 200 | Jandt et al 2015 |
| | HSP 90A R | GTCAATTTGGCGTTGGTTTCTA | | | |

**Supplementary Table 4:** The results of pairwise PERMANOVA analyses investigating differences in the pan microbiota of different samples. (SF=Female from Solitary Foundress Nest; MF=Female from Multiple Foundress Nest; UM= Unparasitized Male; PF= Parasitized Female; PM= Parasitized Male)

| **Groups** | ***p* value** |
|---|---|
| SF (gut and carcass combined) – MF (gut and carcass combined) | 0.006 |
| UM (gut and carcass combined) – MF (gut and carcass combined) | 0.005 |
| UM (gut and carcass combined) – SF (gut and carcass combined) | 0.009 |
| Unparasitized gut (SF, MF and UM) – Unparasitized carcass (SF, MF and Unparasitized male) | 0.84 |
| Parasitized_gut (male and female combined) - Parasitized_carcass (male and female combined) | 0.002 |
| SF (gut and carcass combined) – *X. gadagkari* (male and female combined) | 0.004 |
| MF (gut and carcass combined) – *X. gadagkari* (male and female combined) | 0.002 |
| UM (gut and carcass combined) - *X. gadagkari* (male and female combined) | 0.001 |
| PF (gut and carcass combined) - *X. gadagkari* (male and female combined) | 0.004 |
| PM (gut and carcass combined) - *X. gadagkari* (male and female combined) | 0.004 |

**Supplementary Table 5:** The results of pairwise PERMANOVA analyses investigating differences in the core microbiota of different samples. (SF=Female from Solitary Foundress Nest; MF=Female from Multiple Foundress Nest; UM= Unparasitized Male; PF= Parasitized Female; PM= Parasitized Male)

| **Groups** | ***p* value** |
| --- | --- |
| SF (gut and carcass combined) – MF (gut and carcass combined) | 0.015 |
| UM (gut and carcass combined) – MF (gut and carcass combined) | 0.011 |
| UM (gut and carcass combined) – SF (gut and carcass combined) | 0.013 |
| SF (gut and carcass combined) – *X. gadagkari* (male and female combined) | 0.004 |
| MF (gut and carcass combined) – *X. gadagkari* (male and female combined) | 0.001 |
| UM (gut and carcass combined) - *X. gadagkari* (male and female combined) | 0.011 |
| PF (gut and carcass combined) - *X. gadagkari* (male and female combined) | 0.003 |
| PM (gut and carcass combined) - *X. gadagkari* (male and female combined) | 0.005 |

**Supplementary Table 6:** List of bacterial OTUs found across all samples with their number of reads. OTUs with >= 0.3% abundance are presented here. (SF=Female from Solitary Foundress Nest; MF=Female from Multiple Foundress Nest; UM= Unparasitized Male; PF= Parasitized Female; PM= Parasitized Male)

| ***OTUs*** | **SF gut** | **SF carcass** | **SF Larva** | **MF gut** | **MF carcass** | **UM gut** | **UM carcass** | **PF gut** | **PF carcass** | **PM gut** | **PM carcass** | ***X. gadagkari* Male** | ***X. gadagkari* Female** |
|---|---|---|---|---|---|---|---|---|---|---|---|---|---|
| *Achromobacter* | 0 | 3 | 2 | 3 | 16 | 3 | 0 | 1 | 6 | 2 | 0 | 1 | 1 |
| *Acidisoma* | 1 | 1 | 0 | 41 | 0 | 3 | 0 | 0 | 1 | 0 | 0 | 0 | 0 |
| *Acinetobacter* | 6 | 25 | 46 | 12 | 97 | 42 | 7 | 20 | 22 | 21 | 3 | 11 | 6 |
| *Apilactobacillus* | 2 | 2 | 5 | 2 | 8 | 39 | 4 | 281 | 68 | 90 | 10 | 0 | 0 |
| *Asaia* | 0 | 0 | 0 | 5 | 5 | 9 | 1 | 1 | 0 | 1 | 1 | 0 | 0 |
| *Bacillus* | 33 | 17 | 9 | 1125 | 72 | 28 | 4 | 8 | 3 | 7 | 3 | 1 | 1 |
| *Bartonella* | 0 | 1 | 2 | 0 | 2 | 11 | 13 | 5 | 25 | 58 | 86 | 0 | 1 |
| *Bifidobacterium* | 0 | 0 | 0 | 5 | 29 | 0 | 1 | 1 | 1 | 4 | 1 | 1 | 1 |
| *Bombella* | 0 | 0 | 0 | 0 | 8 | 0 | 0 | 0 | 0 | 20 | 0 | 0 | 0 |
| *Bombilactobacillus* | 235 | 1482 | 7 | 8 | 32 | 70 | 75 | 40 | 307 | 103 | 99 | 0 | 0 |
| *Bradyrhizobium* | 0 | 4 | 1 | 23 | 11 | 3 | 0 | 1 | 2 | 0 | 0 | 2 | 1 |
| *Brevibacillus* | 0 | 0 | 1 | 2 | 2 | 0 | 2 | 2 | 0 | 0 | 0 | 0 | 0 |
| *Brevundimonas* | 3 | 4 | 0 | 2 | 19 | 1 | 0 | 1 | 5 | 0 | 1 | 3 | 0 |
| *Burkholderia-Caballeronia-Paraburkholderia* | 1 | 0 | 0 | 5 | 2 | 11 | 0 | 15 | 6 | 5 | 5 | 0 | 0 |
| *Buttiauxella* | 0 | 0 | 1 | 0 | 1 | 5 | 0 | 1 | 0 | 0 | 0 | 0 | 0 |
| *Citrobacter* | 1 | 16 | 0 | 1 | 3 | 2 | 0 | 0 | 2 | 1 | 1 | 0 | 0 |
| *Corynebacterium* | 0 | 3 | 2 | 4 | 12 | 6 | 1 | 2 | 2 | 0 | 41 | 3 | 0 |
| *Desulfohalotomaculum* | 1 | 0 | 0 | 28 | 0 | 0 | 2 | 0 | 1 | 0 | 0 | 0 | 0 |
| *Devosia* | 0 | 0 | 3 | 0 | 5 | 3 | 3 | 0 | 0 | 1 | 0 | 1 | 1 |
| *Enterobacter* | 27 | 15 | 11 | 24 | 33 | 59 | 1 | 7 | 3 | 9 | 2 | 0 | 0 |
| *Enterococcus* | 0 | 0 | 3 | 0 | 5 | 6 | 1 | 2 | 1 | 0 | 1 | 4 | 1 |
| *Erwinia* | 1 | 0 | 7 | 0 | 2 | 3 | 0 | 0 | 0 | 1 | 0 | 1 | 0 |
| *Escherichia* | 2 | 3 | 0 | 0 | 12 | 0 | 3 | 0 | 0 | 0 | 0 | 0 | 1 |
| *Escherichia-Shigella* | 101 | 180 | 4 | 6 | 429 | 4 | 152 | 0 | 2 | 0 | 2 | 26 | 4 |

| | | | | | | | | | | | | | |
|---|---|---|---|---|---|---|---|---|---|---|---|---|---|
| *Fictibacillus* | 1 | 0 | 7 | 3 | 2 | 0 | 7 | 2 | 1 | 0 | 2 | 0 | 0 |
| *Gluconobacter* | 0 | 0 | 1 | 1 | 20 | 3 | 0 | 1 | 1 | 0 | 2 | 0 | 0 |
| *Halomonas* | 0 | 0 | 0 | 0 | 0 | 3 | 0 | 0 | 0 | 0 | 0 | 0 | 0 |
| *Klebsiella* | 7 | 5 | 20 | 7 | 9 | 10 | 1 | 0 | 0 | 0 | 1 | 2 | 1 |
| *Kluyvera* | 0 | 9 | 0 | 0 | 0 | 3 | 1 | 2 | 0 | 0 | 1 | 0 | 0 |
| *Kosakonia* | 0 | 0 | 1 | 0 | 1 | 7 | 0 | 0 | 0 | 0 | 0 | 0 | 0 |
| *Lactiplantibacillus* | 1 | 4 | 11 | 11 | 14 | 9 | 1 | 0 | 0 | 0 | 0 | 0 | 0 |
| *Lactobacillus* | 165 | 974 | 13 | 10 | 41 | 56 | 53 | 112 | 206 | 78 | 97 | 3 | 5 |
| *Lactococcus* | 345 | 249 | 6 | 197 | 1032 | 30 | 2 | 200 | 84 | 487 | 350 | 2 | 0 |
| *Mesorhizobium* | 0 | 1 | 5 | 6 | 12 | 0 | 1 | 0 | 2 | 0 | 1 | 1 | 0 |
| *Micrococcus* | 0 | 3 | 2 | 6 | 14 | 5 | 1 | 0 | 1 | 1 | 0 | 0 | 0 |
| *Moellerella* | 56 | 307 | 16515 | 2 | 0 | 0 | 0 | 0 | 0 | 0 | 0 | 0 | 0 |
| *Novosphingobium* | 1 | 1 | 3 | 1 | 2 | 3 | 0 | 0 | 0 | 0 | 1 | 0 | 0 |
| *Ochrobactrum* | 2 | 0 | 1 | 4 | 10 | 5 | 4 | 4 | 1 | 3 | 0 | 1 | 1 |
| *Paenibacillus* | 0 | 8 | 2 | 0 | 5 | 2 | 2 | 0 | 1 | 0 | 1 | 0 | 0 |
| *Pantoea* | 1 | 3 | 4 | 2 | 0 | 4 | 4 | 3 | 3 | 1 | 2 | 1 | 0 |
| *Paracoccus* | 0 | 1 | 1 | 9 | 8 | 4 | 1 | 3 | 2 | 2 | 1 | 0 | 0 |
| *Pedomicrobium* | 2 | 2 | 0 | 19 | 3 | 6 | 2 | 4 | 3 | 0 | 0 | 0 | 0 |
| *Pelagibacterium* | 0 | 0 | 0 | 0 | 0 | 0 | 2 | 0 | 0 | 0 | 0 | 0 | 0 |
| *Pelomonas* | 0 | 34 | 1 | 56 | 34 | 1 | 0 | 7 | 1 | 0 | 0 | 4 | 0 |
| *Providencia* | 10 | 21 | 622 | 5 | 9 | 8 | 5 | 2327 | 3690 | 1081 | 416 | 3640 | 2713 |
| *Pseudomonas* | 0 | 27 | 9 | 71 | 89 | 33 | 4 | 15 | 1 | 6 | 4 | 9 | 1 |
| *Pseudoxanthomonas* | 0 | 0 | 2 | 0 | 0 | 149 | 11 | 0 | 1 | 1 | 1 | 0 | 0 |
| *Rothia* | 1 | 5 | 0 | 4 | 7 | 3 | 0 | 0 | 1 | 1 | 0 | 0 | 0 |
| *Salmonella* | 2 | 1 | 10 | 1 | 0 | 4 | 1 | 0 | 0 | 0 | 0 | 0 | 0 |
| *Sediminibacterium* | 1 | 1 | 0 | 16 | 6 | 3 | 1 | 7 | 0 | 1 | 0 | 1 | 0 |
| *Serratia* | 1 | 11 | 0 | 8 | 24 | 2 | 0 | 2 | 4 | 1 | 0 | 1 | 1 |
| *Spiroplasma* | 0 | 8 | 3 | 3213 | 531 | 2 | 0 | 2 | 1 | 0 | 1 | 0 | 0 |
| *Staphylococcus* | 1 | 13 | 15 | 1 | 39 | 9 | 1 | 1 | 5 | 331 | 690 | 5 | 2 |
| *Stenotrophomonas* | 4 | 10 | 34 | 6 | 19 | 8 | 5 | 7 | 21 | 3 | 4 | 2 | 2 |
| *Streptococcus* | 1 | 31 | 7 | 28 | 71 | 10 | 2 | 7 | 6 | 1 | 1 | 2 | 0 |
| *Thioglobaceae* | 0 | 0 | 0 | 0 | 0 | 4 | 0 | 0 | 0 | 0 | 0 | 0 | 0 |
| *Thiomicrorhabdus* | 3 | 1 | 1 | 3 | 1 | 0 | 3 | 0 | 0 | 0 | 1 | 0 | 0 |

| | | | | | | | | | | | | | |
|---|---|---|---|---|---|---|---|---|---|---|---|---|---|
| *Unclassifed_Acetobacteraceae* | 0 | 0 | 0 | 0 | 144 | 0 | 0 | 0 | 0 | 0 | 0 | 0 | 0 |
| *Unclassifed_Caulobacteraceae* | 0 | 0 | 0 | 17 | 8 | 0 | 0 | 0 | 0 | 0 | 0 | 0 | 0 |
| *Unclassified_Acetobacteraceae* | 0 | 1 | 0 | 0 | 0 | 0 | 2 | 1 | 6 | 3 | 7 | 0 | 2 |
| *Unclassified_Blastocatellaceae* | 0 | 0 | 0 | 0 | 0 | 0 | 2 | 1 | 0 | 0 | 0 | 0 | 0 |
| *Unclassified_Caulobacteraceae* | 0 | 3 | 0 | 0 | 0 | 4 | 1 | 2 | 2 | 1 | 0 | 6 | 0 |
| *Unclassified_Moraxellaceae* | 0 | 0 | 0 | 0 | 0 | 0 | 0 | 0 | 0 | 0 | 10 | 0 | 0 |
| *unclassified_Rhizobiaceae* | 0 | 1 | 0 | 0 | 0 | 0 | 0 | 0 | 0 | 0 | 0 | 0 | 0 |
| *Unclassified_Rhodospirillales* | 0 | 0 | 1 | 0 | 0 | 0 | 2 | 0 | 1 | 0 | 0 | 0 | 0 |
| *Unclassified_Thioglobaceae* | 4 | 7 | 1 | 0 | 1 | 0 | 6 | 0 | 13 | 0 | 0 | 0 | 0 |
| *Weissella* | 0 | 22 | 0 | 2 | 2 | 0 | 0 | 0 | 0 | 0 | 0 | 0 | 0 |
| *Wolbachia* | 0 | 2 | 33 | 3 | 6 | 17 | 6 | 6007 | 9938 | 2808 | 1071 | 8960 | 6712 |
| *Xanthomonas* | 0 | 0 | 0 | 0 | 0 | 28 | 3 | 0 | 0 | 0 | 0 | 0 | 0 |
| *Zymobacter* | 0 | 0 | 0 | 0 | 1 | 8 | 0 | 0 | 0 | 0 | 0 | 0 | 0 |

**Supplementary Table 7: Statistical differences in *Wolbachia* density between samples.** One-way ANOVA ($p$ = 0.0024) followed by a multiple comparison test (Tukey's post-hoc test) with a significance level of 0.05. (PF= Parasitized Female; PM= Parasitized Male)

| **Samples** | ***p* value** |
|---|---|
| PF Gut and PF Carcass | 1.0000000 |
| PF Head and PF Carcass | 1.0000000 |
| PM Carcass and PF Carcass | 1.0000000 |
| PM Gut and PF Carcass | 1.0000000 |
| PM Head and PF Carcass | 1.0000000 |
| ***X. gadagkari* Female and PF Carcass** | **0.0220502*** |
| *X. gadagkari* Male and PF Carcass | 0.1632894 |
| PF Head and PF Gut | 0.9999992 |
| PM Carcass and PF Gut | 1.0000000 |
| PM Gut and PF Gut | 1.0000000 |
| PM Head and PF Gut | 1.0000000 |
| ***X. gadagkari* Female and PF Gut** | **0.0280140*** |
| *X. gadagkari* Male and PF Gut | 0.1988003 |
| PM Carcass and PF Head | 1.0000000 |
| PM Gut and PF Head | 1.0000000 |
| PM Head and PF Head | 1.0000000 |
| ***X. gadagkari* Female and PF Head** | **0.0145949*** |
| *X. gadagkari* Male and PF Head | 0.1149058 |
| PM Gut and PM Carcass | 1.0000000 |
| PM Head and PM Carcass | 1.0000000 |
| ***X. gadagkari* Female and PM Carcass** | **0.0228133*** |
| *X. gadagkari* Male and PM Carcass | 0.1679777 |
| PM Head and PM Gut | 1.0000000 |
| ***X. gadagkari* Female and PM Gut** | **0.0226500*** |
| *X. gadagkari* Male and PM Gut | 0.1669781 |
| ***X. gadagkari* Female and PM Head** | **0.0191720*** |
| *X. gadagkari* Male and PM Head | 0.1451902 |
| *X. gadagkari* Male and *X. gadagkari* Female | 0.9877693 |

**Supplementary Table 8:** Bray-Curtis Dissimlarity matrix for all samples. (SF=Female from Solitary Foundress Nest; MF=Female from Multiple Foundress Nest; UM= Unparasitized Male; PF= Parasitized Female; PM= Parasitized Male; XF= *X. gadagkari* Female; XM= *X. gadagkari* Male)

| | **UM Gut** | **UM Carcass** | **MF Gut** | **MF Carcass** | **PF Gut** | **PF Carcass** | **PM Gut** | **PM Carcass** | **SF Larva** | **SF Carcass** | **SF Gut** | **XF** |
|---|---|---|---|---|---|---|---|---|---|---|---|---|
| **UM Carcass** | 0.62 | | | | | | | | | | | |
| **MF Gut** | 0.91 | 0.97 | | | | | | | | | | |
| **MF Carcass** | 0.79 | 0.82 | 0.72 | | | | | | | | | |
| **PF Gut** | 0.94 | 0.97 | 0.95 | 0.93 | | | | | | | | |
| **PF Carcass** | 0.96 | 0.97 | 0.98 | 0.97 | 0.26 | | | | | | | |
| **PM gut** | 0.89 | 0.93 | 0.94 | 0.83 | 0.39 | 0.56 | | | | | | |
| **PM Carcass** | 0.86 | 0.88 | 0.93 | 0.82 | 0.69 | 0.79 | 0.39 | | | | | |
| **SF Larva** | 0.98 | 0.99 | 0.99 | 0.98 | 0.94 | 0.95 | 0.93 | 0.95 | | | | |
| **SF Carcass** | 0.84 | 0.82 | 0.90 | 0.77 | 0.93 | 0.92 | 0.88 | 0.84 | 0.95 | | | |
| **SF Gut** | 0.71 | 0.62 | 0.89 | 0.68 | 0.92 | 0.93 | 0.81 | 0.71 | 0.98 | 0.61 | | |
| **XF** | 0.99 | 0.99 | 1.00 | 0.99 | 0.10 | 0.21 | 0.46 | 0.76 | 0.95 | 0.99 | 0.99 | |
| **XM** | 0.99 | 0.99 | 0.99 | 0.99 | 0.23 | 0.07 | 0.56 | 0.81 | 0.95 | 0.99 | 0.99 | 0.15 |

**Supplementary Table 9:** List of samples used in this study.

| Organism | Order and Family | Tissue | Source/Reference |
|---|---|---|---|
| *Vespa velutina* | Hymenoptera: Vespidae | Gut | Cini et al., 2020 |
| *Vespula pensylvanica* | Hymenoptera: Vespidae | Whole-Body | Rothman et al., 2021 |
| *Apis mellifera* | Hymenoptera: | Gut | Tola et al., 2020 |
| *Formica exsecta* | Hymenoptera: Formicidae | Whole-Body | Jackson et al., 2023 |
| *Formica lemani* | Hymenoptera: Formicidae | Whole-Body | Jackson et al., 2023 |
| *Dipeterophagus daci* | Strepsiptera: Halictophagidae | Whole body | Towett-kirui et al., 2023 |

**Supplementary Figure 1:** Heatmap of bacterial OTUs that accounted for equal or greater than 0.3% abundance in one or more samples. (SF=Female from Solitary Foundress Nest; MF=Female from Multiple Foundress Nest; UM= Unparasitized Male; PF= Parasitized Female; PM= Parasitized Male; G= Gut; C= Carcass)

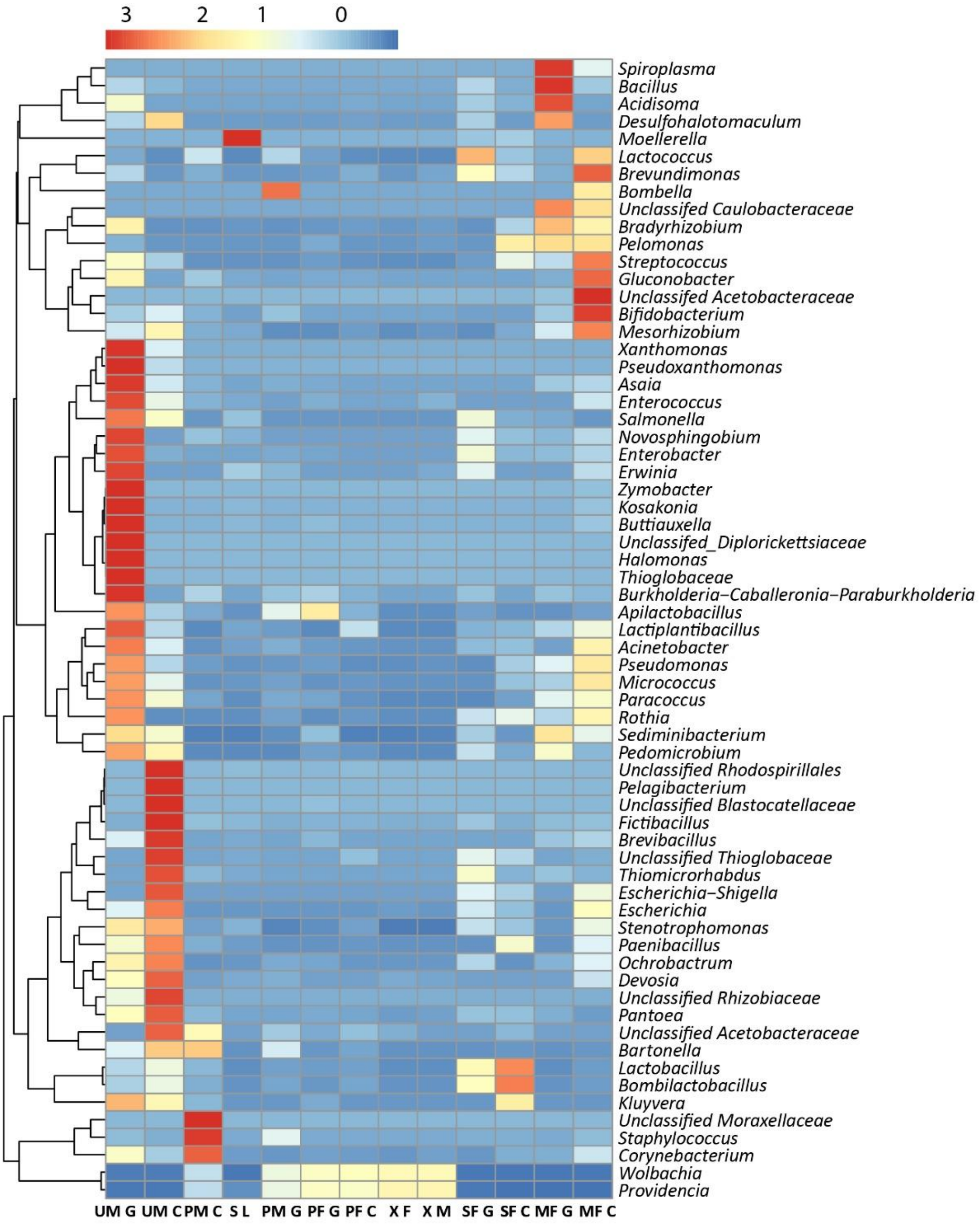

**Supplementary Figure 2: Rarecurve Analysis.** Rarefaction curve showing the number of OTUs on Y-axis and number of identified bacterial reads on X-axis for *P. wattii* and *X. gadagkari* samples.

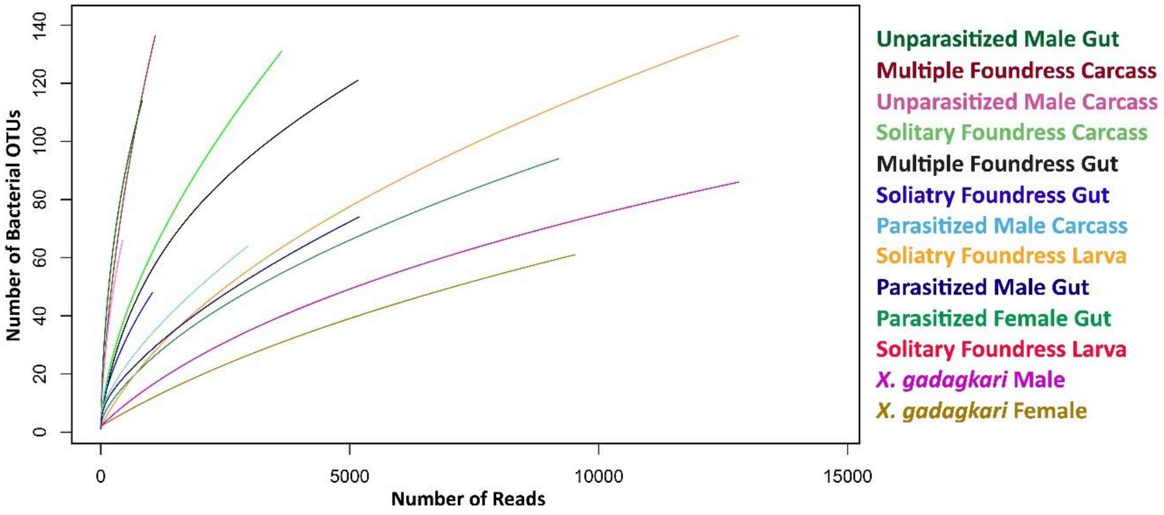

**Supplementary Figure 3: Core Microbiome of *P. wattii*. a) Venn diagram showing the number of unique and shared bacterial genera among unparasitized *P. wattii*** (SF= Female from Solitary foundress nest; MF= Female from multiple foundress nest; UM= Unparasitized male)**.** 16 bacteria constitute the core microbiota of unparasitized *P. wattii,* including *Sediminibacterium, Escherichia-Shigella, Pedomicrobium, Providencia, Stenotrophomonas, Lactiplantibacillus, Staphylococcus, Klebsiella, Streptococcus, Bacillus, Lactococcus, Apilactobacillus, Acinetobacter, Lactobacillus*, *Enterobacter* and *Bombilactobacillus.* **b) Venn diagram showing the number of unique and shared bacterial genera between unparasitized *P. wattii* and *X. gadagkari*.** 8 bacteria are shared between the core microbiota of unparasitized *P. wattii* and *X. gadagkari* including *Escherichia-Shigella*, *Stenotrophomonas*, *Bacillus, Acinetobacter, Lactobacillus, Staphylococcus, Providencia*, and *Klebsiella*.

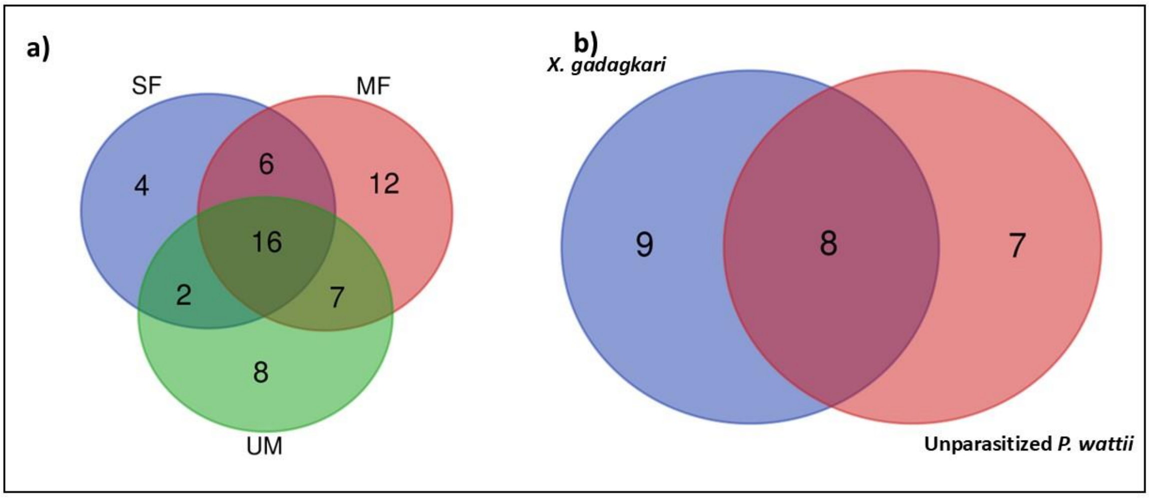

**Supplementary Figure 4:** Relative *Wolbachia* density was measured in *P. wattii* and *X. gadagkari* samples. Three biological replicates were tested for each sample. Each bar represents an average of three biological replicates. Tukey's HSD post hoc test was used for multiple comparisons at the $p < 0.05$ level. (SF=Female from Solitary Foundress Nest; UM= Unparasitized Male; PF= Parasitized Female; PM= Parasitized Male)

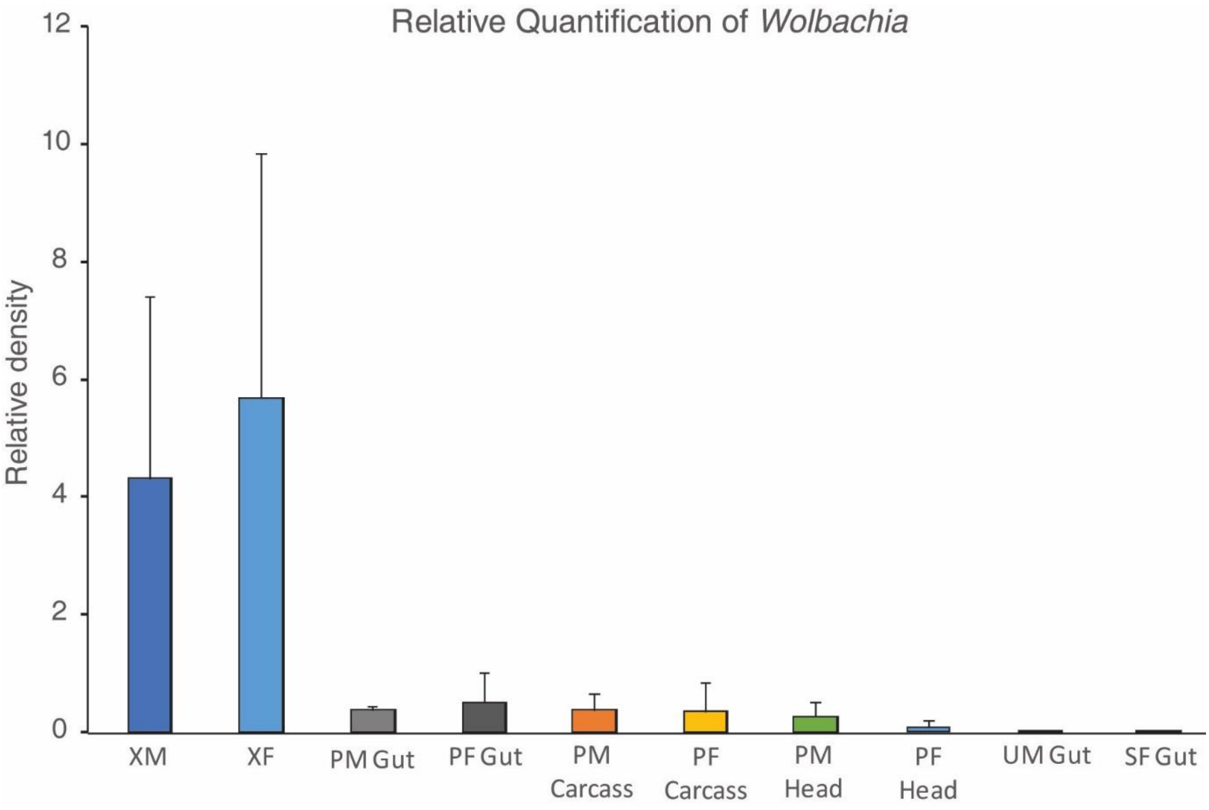
